# A blueprint for human whole-cell modeling

**DOI:** 10.1101/198044

**Authors:** Balázs Szigeti, Yosef D. Roth, John A. P. Sekar, Arthur P. Goldberg, Saahith C. Pochiraju, Jonathan R. Karr

## Abstract

Whole-cell models of human cells are a central goal of systems biology. Such models could help researchers understand cell biology and help physicians treat disease. Despite significant challenges, we believe that human whole-cell models are rapidly becoming feasible. To develop a plan for achieving human whole-cell models, we analyzed the existing models of individual cellular pathways, surveyed the biomodeling community, and reflected on our experience developing whole-cell models of bacteria. Based on these analyses, we propose a plan for a project, termed the *Human Whole-Cell Modeling Project*, to achieve human whole-cell models. The foundations of the plan include technology development, standards development, and interdisciplinary collaboration.

## Introduction

Over the past 25 years, researchers have built numerous models of individual cellular pathways. However, to comprehensively understand cells, we must build whole-cell (WC) models that represent multiple pathways and the interactions among them [1, 2]. WC models of human cells could transform medicine by helping researchers gain insights into complex phenotypes such as development and by helping physicians personalize therapy [3–5].

Recently, we and others demonstrated that WC models are feasible by developing the first model that represents the function of every characterized gene in a bacterium [6]. However, it took over took over ten person-years to build the model, in part because there are few data sources, model design tools, model description formats, and simulators suitable for WC modeling [2, 7–9]

Here, we outline a plan for scaling up from models of individual human pathways to WC models of entire human cells. To develop the plan, we analyzed existing models and modeling methods; surveyed the community to determine the major bottlenecks of biomodeling and new tools that are needed to advance biomodeling; and reflected on our experience developing WC models of bacteria. Lastly, we propose a plan for a project, called the *Human Whole-Cell Modeling Project*, to achieve human WC models.

## Building blocks for WC models: models of individual pathways

A wide range of models of individual pathways could be leveraged to build human WC models (Figure 1, Table S1). Notably, there are several detailed kinetic models of individual signaling pathways including the JAK/STAT, NF-κB, p53, and TGF β pathways [10], as well as detailed kinetic models of the cell cycle [11], circadian rhythms [12], and electrical signaling [13]. Most of these models are represented as reaction networks that can be simulated by integrating ordinary differential equations (ODEs). However, some of the largest of these models must be represented using rules [14] and simulated using network-free methods [15] to efficiently manage their combinatorial complexity.

**Figure 1.**
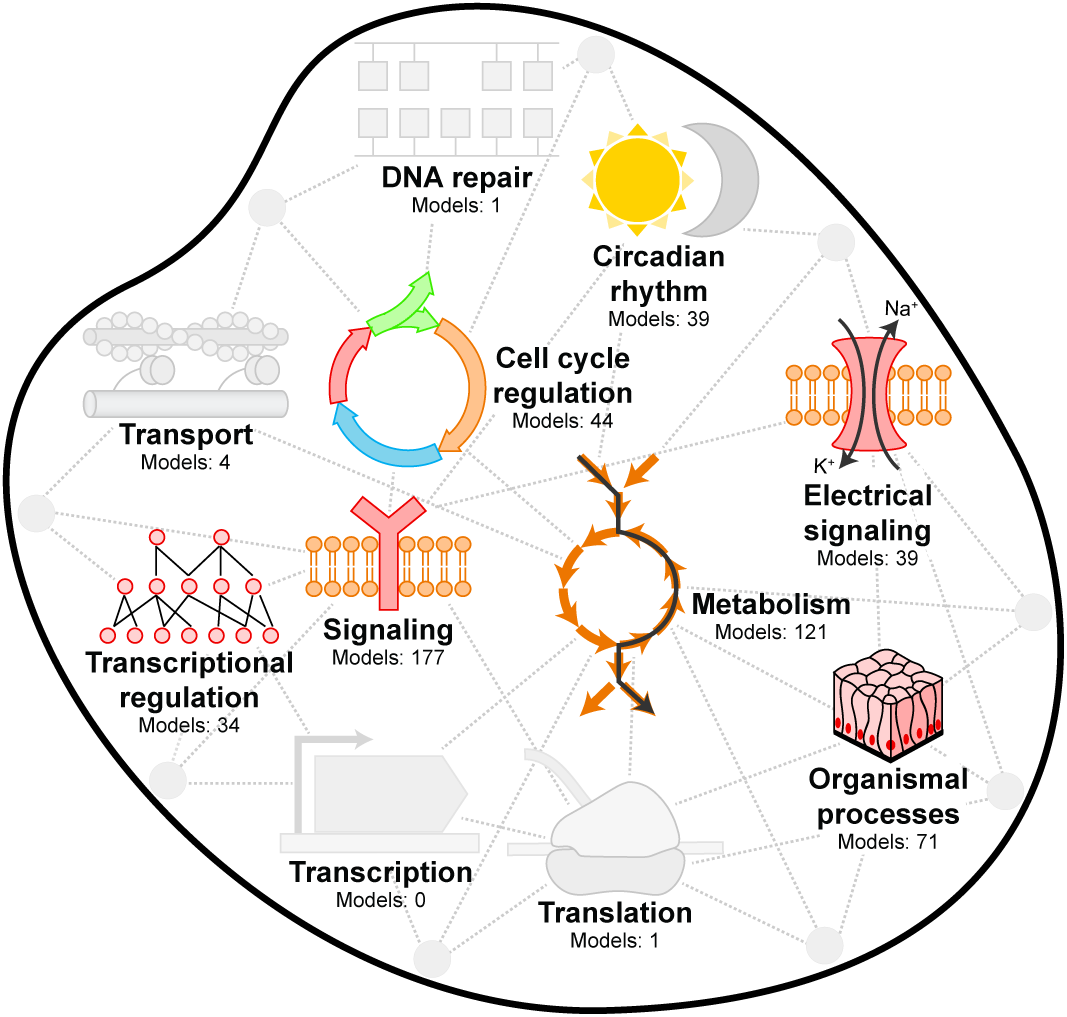
WC models can be built by leveraging existing models of well-studied processes (colors) and developing new models of other processes (gray).

There are also several detailed models of human metabolism [16–18] and transcriptional regulation [19]. The largest metabolism models use flux balance analysis (FBA) to predict steady-state reaction fluxes [20]. The largest transcriptional regulation models use Boolean networks to describe the regulatory logic of each gene and transcription factor [21].

In addition, there are a small number of models of other pathways such as apoptosis and differentiation, as well as a few models of multicellular processes such as tissue patterning and infection[22]

Despite this progress, many pathways have not been modeled at the scale of entire cells, including several well-studied pathways. For example, although we have extensive knowledge of the mutations responsible for cancer, we have few models of DNA repair; although we have extensive structural and catalytic information about RNA modification, we have few kinetic models of RNA modification; and although we have detailed atomistic models of protein folding, we have few cell-scale models of chaperone-mediated folding.

## Pioneering efforts to build integrative models of multiple pathways

Since 1999 when Tomita *et al.* reported one of the first integrative models of *M. genitalium* [23], researchers have been trying to build increasingly comprehensive models of multiple pathways. This has led to models of *Escherichia coli* and *Saccharomyces cerevisiae* which describe their metabolism and transcriptional regulation [24, 25]; their metabolism, signaling, and transcriptional regulation [26–28]; and their metabolism and RNA and protein synthesis and degradation [29]. The most comprehensive cell model to date represents 28 pathways of *Mycoplasma genitalium* [6]. Table S2 summarizes several recently published and proposed integrative models, including several integrative models of human cells [30–34].

One of the main challenges to modeling multiple pathways is that the depth of our knowledge, and consequently the detail of our models, varies across cellular pathways. For example, although we have detailed kinetic information about several signaling pathways, we have little kinetic data about metabolism. Similarly, although we have extensive information about the participants in thousands of metabolic reactions, we have incomplete knowledge about the targets of each signaling mediator. Consequently, metabolic models are often simulated using FBA [20], whereas signaling models are often simulated using ODEs and stochastic simulation algorithms. As a result, most models of multiple pathways combine multiple simulation formalisms such as Boolean networks, FBA, ODEs, and stochastic simulation. However, few tools support the construction and simulation of such multi-algorithmic models.

## Bottlenecks to WC modeling

To help us formulate a plan for progressing to WC models of human cells, we invited the community to participate in a survey of the limitations of our modeling methods and tools. As detailed in Note S1A, we asked the community three sets of questions: (a) multiple response questions about their sector, field, and country; (b) multiple response questions about their research goals, methods, and the challenges to their research; and (c) open-ended questions about new goals, methods, tools, and resources that could advance biomodeling.

As detailed in Note S1B, we recruited a broad range of scientists to participate in the survey by identifying scientists from the Europe PMC database [35]; the NSF award database (URL: https://www.nsf.gov/awardsearch); community, conference, and university websites; and our personal contacts. In total, we invited 542 scientists from a wide range of sectors and countries (Figures S1–S2). We also sent survey invitations to three community email lists, and we encouraged scientists to share our invitation with their colleagues.

As summarized in Figures S3–S7, 214 scientists completed the survey. 94% of the respondents are employed in academia and 6% are employed in industry. The respondents spanned a wide range of fields including systems biology (41%), mathematical biology (21%), multiscale modeling (15%), and bioinformatics (12%) (Figure 2A); the respondents had a wide range of expertise including modeling and simulation (85%); methods, software, and standards development (57%), and experimentation (15%); the respondents focused on a wide range of biological systems including signaling (34%), metabolism (38%), and transcriptional regulation (23%); and the respondents spanned a wide geographical range including Europe (22%), the USA (21%), and Asia (6%)

**Figure 2.**
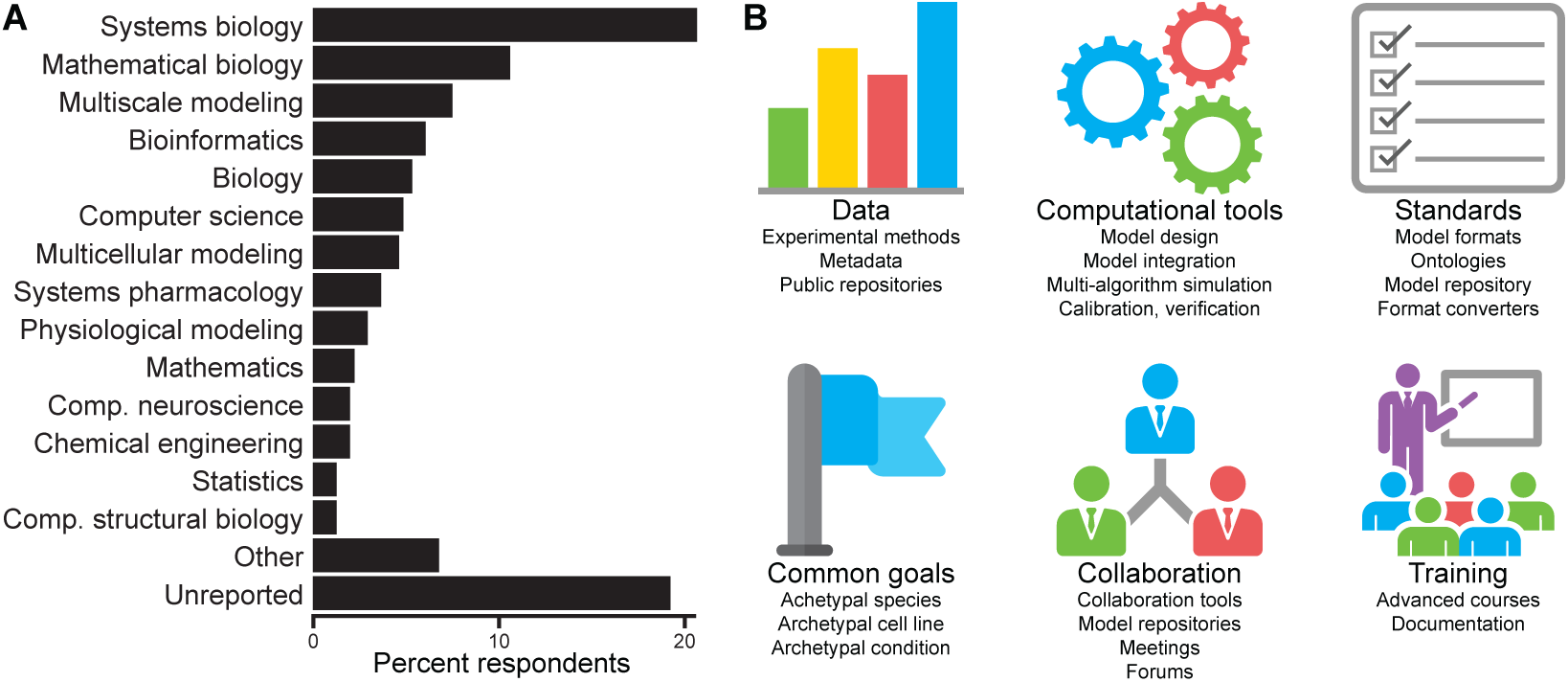
A survey of the biomodeling community indicated that WC models can be achieved by collaborating on a common goal, developing scalable modeling tools, sharing data and models, adopting modeling standards, and training new researchers. (**A**) Fields of the 214 respondents. (**B**) Major bottlenecks to biomodeling and the major methods, tools, and resources needed to advance biomodeling reported by the respondents.

To our knowledge, we conducted the first survey of the biomodeling community since a 2013 survey of European systems biologists conducted by the Infrastructure for Systems Biology Europe [36] and a 2015 survey of Canadian bioinformaticians and computational biologists (URL: https://bioinformatics.ca/bcb/survey).

Here, we summarize the key bottlenecks to biomodeling reported by the community (Figure 2B, Figures S8–S21), and use our WC modeling experience to analyze their implications for WC modeling. Although the survey focused on biomodeling in general, we believe that the reported bottlenecks are also major roadblocks to WC modeling.

### Inadequate experimental methods and data repositories

The most commonly reported bottleneck to biomodeling, by far, was the lack of experimental data for modeling (82%). In addition, data aggregation was the most commonly reported time-consuming aspect of biomodeling (53%). Furthermore, the most commonly reported data sources for biomodeling were articles (83%) and supplementary materials (71%), which, in our experience often require extensive manual effort [6].

Because WC models will require comprehensive experimental data about the structure, abundance, interactions, and dynamics of molecular species [2], we believe that these bottlenecks will be especially acute for WC modeling. Based on these findings and our experience, we believe that new measurement methods, data repositories, and data aggregation tools are needed for WC modeling: (a) improved proteome-wide methods for measuring protein abundances would facilitate more accurate models of many pathways; (b) improved metabolome-wide methods for measuring metabolite concentrations would enable more accurate models of metabolism; (c) new single-cell measurement methods would facilitate more accurate models of the phenotypic variation of single cells; (d) a new central data repository that uses consistent representations, identifiers, and units would accelerate data aggregation [37]; and (e) new tools for searching this repository would help researchers identify relevant data for WC modeling, including data from related organisms and environments.

### Incompatible models

A second commonly reported bottleneck to biomodeling was the incompatibility among models (30%). In our and others’ experiences, there are several reasons why models are often incompatible [38–41]: (a) many models are difficult to obtain or are never published. For example, only 36% of survey respondents reported that they deposit models to repositories such as BioModels (b) Models are often described using inconsistent or idiosyncratic formats. For example, only 55% of respondents reported that they use the Systems Biology Markup Language [42] (SBML). (c) The biological meaning of each model component is rarely annotated. (d) Models are often described using inconsistent identifiers and units. (e) Few models are described with consistent granularity. For example, some models represent individual reactions, whereas other models describe lumped reactions. (f) Models of separate pathways rarely have clearly and consistently defined interfaces. For example, models of separate signaling pathways often share metabolites and proteins. (g) Few models are based on consistent assumptions. For example, signaling models often assume that gene expression is constant, whereas models of transcriptional regulation often predict the dynamics of gene expression.

To enable WC modeling, we must resolve these incompatibilities because WC models will likely be built by combining separately developed models of individual pathways. To increase the compatibility of models, we believe that we should (a) develop better formats for describing the semantic meaning, data sources and the assumptions used to build models; (b) develop better tools for annotating models; and (c) require researchers to use standard formats and model repositories.

### Inadequate tools for designing, calibrating, and validating models

Many scientists reported that the lack of adequate model design (15%), calibration (21%), and validation (31%) tools are also significant bottlenecks to biomodeling. To overcome these bottlenecks, we believe that the field should develop software tools for (a) designing models directly from experimental data, (b) efficiently calibrating high-dimensional models, and (c) systematically checking that models are consistent with experimental data.

### Inadequate model formats

14% of survey respondents also reported that the lack of adequate model formats is a bottleneck to biomodeling. Most survey respondents reported that they describe models using reaction network formats such as SBML [42] (54%) or rule-based formats such as BioNetGen [14] (7%). In our experience, no existing format is well-suited to describing WC models because no existing format can represent (a) the combinatorial complexity of pathways such as transcription elongation which involve billions of sequence-based reactions; (b) the multiple scales that must be represented by WC models such as the sequence of each protein, the subunit composition of each complex, and the DNA binding of each complex; and (c) models that are composed of multiple mathematically-distinct submodels [8].

To enable human WC models, we must develop a format which is capable of (a) describing models in terms of high-level biological constructs such as DNA, RNA, and proteins; (b) representing all of the types of combinatorial complexity that arise in molecular biology including the combinatorial number of possible interactions of sites among within macromolecules and the combinatorial number of RNA and proteins that can arise from processes such as splicing, editing, and mutations; (c) explicitly representing the data and assumptions used to build models; and (d) representing the biological meaning of models. Furthermore, to clearly represent models, we believe that this format should be based on rules.

## Realizing WC models of human cells

Despite these challenges, we believe that we should begin to develop a plan for achieving human WC models because human WC models are rapidly becoming feasible thanks to new modeling and experimental technologies [2, 14, 15, 43–47] and because human WC models have great potential to transform science and medicine. Below, we first propose several guiding principles for a human WC modeling project, and then we propose a concrete, three-phase plan for achieving human WC models.

### Guiding principles for human WC modeling

Based on the literature, our survey, and our modeling experience, we propose eight guiding principles for a human WC modeling project. (a) The project should develop technologies for scalably building, calibrating, simulating, and validating WC models. (b) To facilitate collaboration, the project should develop standards for describing WC models and standard protocols for validating and merging model components. (c) The project should build models collaboratively by partitioning cell biology into distinct pathways, outlining the interfaces among these pathways, tasking different researchers to build and validate composable submodels of each pathway, and merging the submodels into a single model. (e) The project should build human WC models in close coordination with extensive experimental characterization of human cells. (e) The project should form an interdisciplinary community of modelers, experimentalists, computer scientists, and engineers, as well as research sponsors to achieve WC models. (f) The project should focus on a single cell line that is easy to culture, well-characterized, karyotypically and phenotypically “normal” genomically stable and relevant to a wide range of basic science and medicine such as the H1 human embryonic stem cell (hESC) line. (g) The methods and models developed by the project should continuously evolve as we acquire new information. This should include how we partition cells into pathways and the interfaces that we define among the pathways. (h) Based on lessons learned from other “big science” projects, the project should delineate clear subgoals, clearly define the responsibilities of each researcher, and freely exchange information [48, 49].

### A plan for the Human Whole-Cell Modeling Project

More concretely, we propose the following three-phase plan for achieving the first WC model of a human cell (Figure 3). To focus the community’s effort, the plan focuses on modeling H1-hESCs, which we believe are an ideal testbed for human WC modeling. However, the methods and tools developed by the project would also be applicable to any organism and the H1-hESC model could be contextualized with genome-scale data to represent other cell lines, cell types, and individuals.

**Figure 3.**
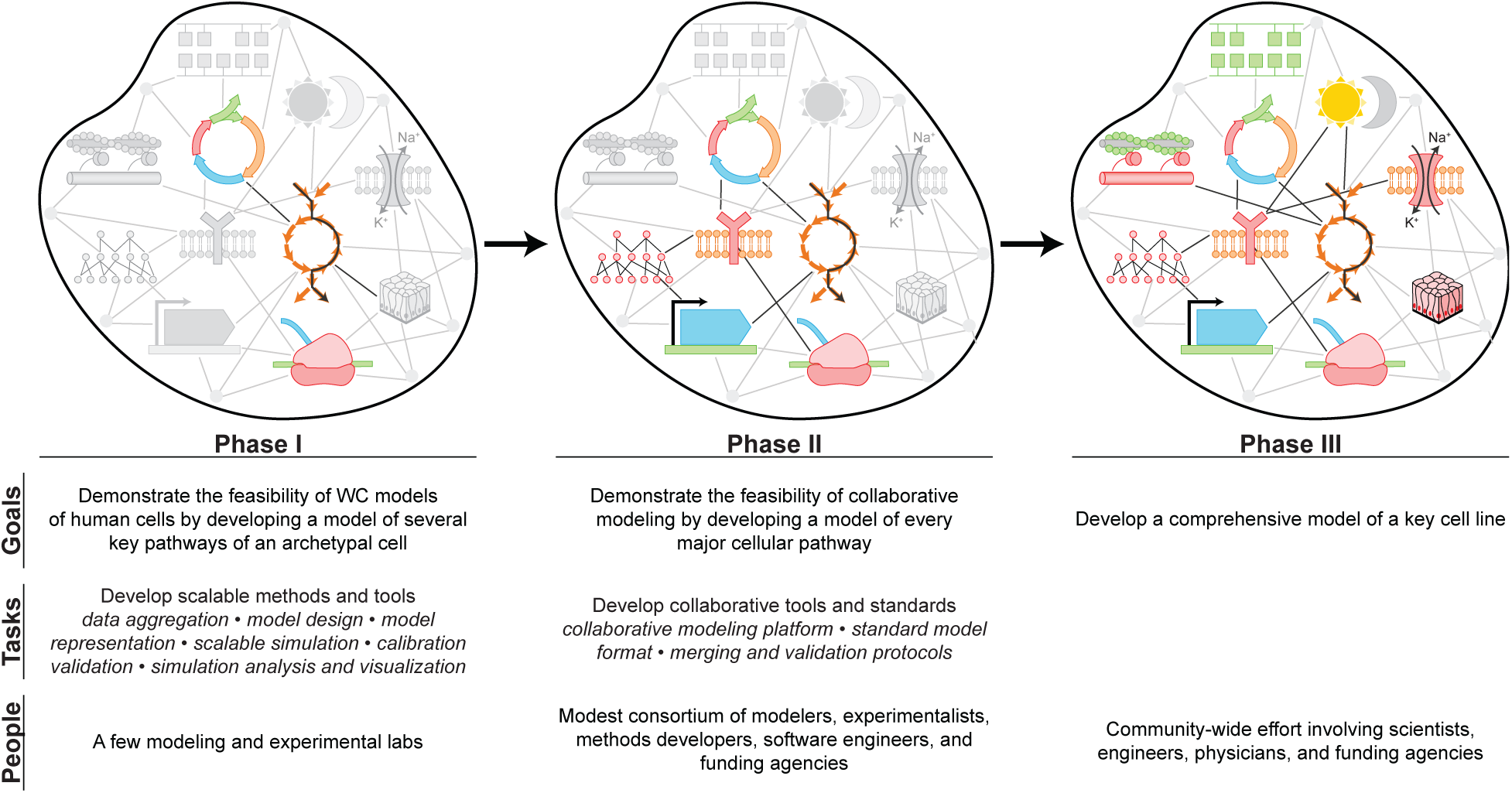
The proposed *Human Whole-Cell Modeling Project* would achieve human WC models in three phases: (1) demonstrating the feasibility of human WC models by developing scalable modeling tools and using them to model several core processes, (2) demonstrating the feasibility of collaborative modeling by developing a collaborative modeling platform and using it to model additional processes, and (3) developing a comprehensive model as a community.

### Phase I: Piloting the core technologies and concepts of human WC modeling

Phase I should demonstrate the feasibility of human WC models by developing the core technologies needed for WC modeling, and using these tools to build a model of a few critical pathways of H1-hESCs. First, we should develop tools for aggregating the data needed for WC modeling, tools for designing models directly from data, a rule-based language for describing models, tools for quickly simulating multi-algorithmic models, tools for efficiently calibrating and validating high-dimensional models, and tools for visualizing and analyzing high-dimensional simulation results. Second, a small group of researchers should use these tools and public data to build a model of the core pathways of H1-hESCs including several key signal transduction pathways, metabolism, DNA replication, transcription, translation, and RNA and protein degradation. In Phase I, we should also begin to form a WC modeling community by organizing meetings and discussing WC modeling standards.

### Phase II: Piloting collaborative WC modeling

Phase II should focus on demonstrating the feasibility of collaborative WC modeling by developing collaborative modeling tools, and using them to expand the H1-hESC model begun in Phase I. First, we should combine the technologies developed in Phase I into a collaborative web-based WC modeling platform to enable multiple experts to build models together. Second, we should develop standards for describing, validating, and merging submodels. Third, we should expand the H1-hESC model developed in Phase I by forming a small consortium of modelers and experimentalists, partitioning cell biology into distinct pathways, outlining the interfaces among these pathways, and tasking individual researchers with modeling additional pathways such as cell cycle regulation, DNA repair, and cell division. Fourth, we should extensively validate the combined model. In Phase II, we should also continue to develop the fundamental technologies needed for WC modeling and continue to build a WC community.

### Phase III: Community modeling and model validation

Phase III should produce the first comprehensive WC model of a human cell. First, we should assemble a large community of modelers and experimentalists and train them to use the platform developed in Phases I and II. Second, we should task individual researchers with building models of individual pathways and merging them into the global H1-hESC model. Third, we should continue to validate the combined model. Fourth, we should use the model to generate testable hypotheses to discover new biology, new disease mechanisms, and new drug targets. Fifth, we should also begin to develop methods for contextualizing the H1-hESC model to represent other cell lines, cell types, and individuals. In addition, we should continue develop the core technologies and standards needed for WC modeling, continue to refine the partitioning of cells into pathways, and continue to refine the interfaces among the pathways.

## Conclusions

In summary, we believe that human WC models have great potential to transform basic science and medicine. Furthermore, we believe that human WC models are rapidly becoming feasible thanks to advances in measurement and modeling technology.

To advance human WC modeling, we summarized the current state of human cell modeling by reviewing the literature, identified the major bottlenecks to WC modeling by surveying the community, and proposed a plan for a project, termed the *Human Whole-Cell Modeling Project*, to achieve human WC models. The cornerstones of our plan include developing computational technologies for scalably building and simulating models, developing standard protocols and formats to facilitate collaboration, building models collaboratively as a community, and focusing on a single cell line. We are excited to embark on this ambitious plan, and hope that you join our efforts to build human WC models!

## Acknowledgements

We thank Yin Hoon Chew for critical feedback on the manuscript and we thank Michael Hucka, Derek Macklin, and Pedro Mendes for critical feedback on the survey questions. This work was supported by a National Institute of Health MIRA award [grant number 1 R35 GM 119771-01]; a National Science Foundation INSPIRE award [grant number 1649014]; and the National Science Foundation / ERASynBio [grant numbers 1548123, 335672].

Papers of particular interest, published within the period of review, have been highlighted as:

*•* of special interest

*••* of outstanding interest

